# Sex-Specific Vulnerability to Radiofrequency Electromagnetic Radiation-Induced Reproductive and Neurological Impairment in Mice

**DOI:** 10.64898/2026.03.06.710073

**Authors:** Kai Zhu, Feilong Li, Zhiyong Liu, Jiaming Guo, Xiwen Yang, Chanyi Li, Jiaming Shen, Linhui Wang, Hongli Yan

**Affiliations:** Center of Reproductive Medicine, Changhai Hospital, Naval Medical University, Shanghai, China; Urology Department, Changhai Hospital, Naval Medical University, Shanghai, China; Department of Radiation Medicine, College of Naval Medicine, Naval Medical University, Shanghai, China; First Affiliated Hospital, Henan University, Henan, China

**Keywords:** electromagnetic radiation, reproduction, testis, hippocampus, neuron, biomarkers

## Abstract

Electromagnetic radiation (EMR) constitutes a pervasive environmental stressor, yet the sex-specific vulnerabilities to its biological effects remain inadequately characterized. This study investigated the differential impacts of 3.2 GHz pulsed EMR on reproductive and neurological functions in male and female mice following a four-week exposure regimen. We identified a pronounced sexual dimorphism in systemic and organ-specific responses. Male mice exhibited a primary vulnerability in the reproductive axis, manifesting as severe spermatogenic impairment, including depletion of the spermatogonial stem cell pool, disruption of sperm chromatin remodeling, and consequent declines in sperm quality and quantity. In contrast, exposed females showed preserved ovarian function but displayed marked neurobehavioral deficits, including increased anxiety- and depressive-like behaviors, and cognitive impairment, which were correlated with neurodegeneration in the hippocampus and prefrontal cortex. Proteomic analyses revealed distinct sex-specific biomarker profiles, with serum levels of Mdk, histone H1 variants (H1-1, H1-4, H1-5), Hp1bp3, Ncf2, and Rhoc strongly associated with male testicular dysfunction, while decreased levels of the synaptic motor protein KIF13A in serum and brain tissue were linked to female neurological impairment. Our findings delineate clear sex-divergent pathological pathways that target male spermatogenesis and female neurocognitive function and propose novel circulating protein biomarkers. This work provides a critical foundation for sex-aware risk assessment and the development of targeted preventive strategies against EMR-associated health risks.

## 1. Introduction

The rapid advancement of wireless communication technologies since the 1950s has led to an exponential increase in ambient non-ionizing electromagnetic radiation (EMR). Peak power flux densities now exceed natural background levels by more than 10¹⁸ times [1–3], particularly within the radiofrequency (RF-EMR) range of 300 kHz to 300 GHz [4]. In response to growing public health concerns, the World Health Organization (WHO) has recognized electromagnetic radiation as a significant environmental pollutant, and the International Agency for Research on Cancer (IARC) has classified it as “possibly carcinogenic to humans” (Group 2B) [5]. Upon absorption by biological tissues, RF-EMR induces not only thermal effects but also triggers non-thermal biological responses, including dysregulated cell proliferation, altered gene expression, DNA damage, and oxidative stress. These mechanisms collectively impair sensitive physiological systems, notably reproduction and neurophysiological function, with emerging evidence suggesting these adverse effects may exhibit sex-specific patterns. This underscores the urgent need to elucidate sex-based differences in pathophysiological mechanisms and to identify reliable biomarkers for accurate risk assessment.

Accumulating evidence indicates that RF-EMR adversely affects reproductive system of both men and women; however, sex-specific susceptibility remains insufficiently explored. In males, exposure is associated with impaired spermatogenesis, characterized by reduced sperm quality, decreased spermatogonial populations, elevated apoptosis, and altered testosterone levels, contributing to diminished fertilization capacity and potential embryonic abnormalities [6, 7]. The testis is particularly vulnerable, with non-ionizing radiation provoking dose-dependent oxidative damage [8, 9]. Higher field intensities and pulse frequencies exacerbate oxidative stress, deplete antioxidant defenses, and hinder testicular recovery [10]. In females, RF-EMR compromises ovarian folliculogenesis, reducing primordial, antral, and preovulatory follicle counts alongside decreased corpora lutea and increased atresia [11]. It also disrupts the hypothalamic-pituitary-gonadal axis, altering the secretion of gonadotropins and sex hormones, which disturbs estrous cyclicity, lowers ovulation rates, and leads to subfertility [12, 13].

The central nervous system (CNS) is another primary target due to its electrophysiological properties. Chronic RF-EMR exposure impairs neuronal morphology, neurogenesis, and cognitive-affective processes such as learning, memory, and mood regulation [14, 15]. It also compromises the integrity of the blood-brain barrier (BBB)[16], with studies showing RF-EMR-induced focal albumin leakage and neuronal albumin uptake. This leads to neurodegeneration and altered neurotransmission in critical regions like the hippocampus, hypothalamus, and striatum[17, 18]. Prolonged exposure results in marked hippocampal neurodegeneration, neuronal pyknosis, and vascular congestion, supported by electrophysiological and behavioral evidence of impaired synaptic plasticity [19]. Notably, emerging data suggest a sex-dependent neurovascular vulnerability; for instance, male rats exhibit more pronounced albumin extravasation and cerebrovascular permeability post-exposure than females, implying greater male susceptibility [17, 18]. These findings highlight the necessity for systematic, sex-stratified research to decipher the molecular basis of such differential effects.

Although RF-EMR is established as detrimental to both reproductive and neurological health, it remains unclear whether its impacts differ between sexes in susceptibility, severity, or underlying mechanisms. To address this gap, we exposed male and female mice to 3.2 GHz pulsed RF-EMR for four weeks. This frequency is representative of common Wi-Fi bands and is increasingly relevant in emerging satellite and computational technologies, yet its biological effects remain poorly characterized. Our study provides a systematic, comparative analysis of functional deficits in both reproductive and neural systems between sexes. Furthermore, by conducting integrated proteomic profiling of testicular, brain, and serum samples, we identified circulating protein biomarkers indicative of sex-specific RF-EMR damage. The development of such non-invasive biomarkers may help to enhance the early diagnosis of radiation-induced pathology. Ultimately, this work may inform future risk assessment frameworks and personalized protective strategies against RF-EMR-associated health risks.

## 2. Methods

### 2.1 Experimental animal

Twenty male and twenty female 8-week-old balb/c mice (weighing approximately 25 g) were purchased from GemPharmatech. The mice were housed in a controlled environment with a 12-hour light/12-hour dark cycle and a room temperature of 24°C ± 2°C. The animals were in a stable condition and had free access to standard commercial pellet feed and water. Each cage contained five animals, which were housed in standard ventilated polypropylene cages with dried rice husks as bedding. The 20 randomly selected mice were divided into a control group and an EMR group, with 10 mice in each group. The mice in both experimental groups were continuously exposed to 3.2 GHz (pulsed wave) microwave radiation from 9:00 to 17:00, 8 hours per day, for 4 weeks.

### 2.2 Electromagnetic radiation exposure

The study animals were secured in plastic containers with ventilation holes and positioned beneath a horn antenna orientated for radiation exposure. Each rocket is sufficiently big to permit unrestricted movement of animals and features holes (1 cm in diameter) on the upper and four lateral sides to enhance airflow. Mice in the EMR group were subjected to a 3.2-GHz high-power microwave field for 8 hours every day over a duration of four weeks. The control group experienced identical conditions to the radiation group but was not subjected to microwave exposure. The radiation system comprised a microwave generator, an antechamber, and a control unit. The separation between the antenna and the cage top was 1.2 meters, and microwave pulses were emitted with a pulse duration of 100 microseconds for irradiation. The mean field strength is 90 W/m, and the electromagnetic radiation detector is positioned at the four corners of the enclosure to guarantee the consistency of the radiation field strength received by the animals. The average specific absorption ratio (SAR) of testicular and cerebral tissues was computed utilising the finite-difference time-domain approach via the S4Llite online platform. The formula is SAR = (σ E ^2^)/ 2ρ(E: the electric field strength, σ: electrical conductivity, ρ: tissue density). The average SAR values for brain and testicular tissue in EMR groups were 0.56 W/kg and 0.32 W/kg, respectively. The infrared thermal imaging system monitors real-time fluctuations in temperature and physiological conditions of mice to elucidate the thermal effects during the exposure process.

### 2.3. Computer-Assisted Sperm Analysis (CASA)

Take one side of the cauda epididymis and place it into 1 ml of preheated Hank’s solution at 37°C. Use scissors to shear it 4-5 times to release the sperm. Place the sperm suspension to be tested into a water bath at 37°C and incubate for 10-15 minutes. Take 10 μL of sperm and place it onto a preheated CASA-specific slide and coverslip. Place it under a microscope for real-time observation. Collect sperm motility parameters from 7 different fields of view per sample.

### 2.4 Sex hormone detection

FSH, LH, estradiol, testosterone, inhibin B from serum were quantified using Mouse FSH ELISA Kit (Aviva Systems Biology, OKDD03124), Mouse LH ELISA Kit (Aviva Systems Biology, OKEH00840), Mouse Estradiol ELISA Kit (Aviva Systems Biology, OKEH02540), Mouse Inhibin B ELISA Kit (BiogradeTech, BGT-KET-14309). All ELISAs were run on a 96-well plate and analyzed via colorimetric readout using the Synergy Neo Microplate Reader (BioTek Instruments).

### 2.5 Immunofluorescence (paraffin section)

Place the sections in an oven at 60°C for approximately 15 minutes, then immerse them in xylene, 100% ethanol, 95% ethanol, 85% ethanol, and 70% ethanol for dewaxing and rehydration. Antigen retrieval was performed by boiling in Tris-EDTA buffer (pH=9.0) for 20 mins. Block with 5% donkey serum (Jackson, USA, 017-000-121) at room temperature for 1 hour; incubate with primary antibody overnight at 4°C. Dilute the secondary antibody at a ratio of 1:400 and incubate at room temperature for 1 hour. Scanning and analysis were performed using a digital slide scanner, Nano Zoomer S60. Stained tissue sections from at least three animals per group were analyzed. Immunofluorescence intensity and positive cell counts were quantified using ImageJ software.

### 2.6 Immunofluorometric assay

For TUNEL staining, eight non-overlapping fields of view within the target region were randomly selected for each tissue section. Images for apoptosis quantification were acquired under a 40× objective lens using a slide fluorescence scanner configured for dual-channel imaging: the DAPI channel labeled all cell nuclei, while the FITC channel identified TUNEL^+^ apoptotic cells. Two independent investigators performed blinded counting, defining total cells as DAPI-positive nuclei and apoptotic cells by the presence of irregular punctate or clumped green fluorescence co-localized with DAPI-stained nuclei. The apoptotic rate per field was calculated as (number of apoptotic cells / total cells) × 100%, with the sample apoptotic rate expressed as the mean ± SEM of rates from eight fields. This identical counting methodology was applied to quantify SOX9, STRA8, and NeuN immunofluorescence staining, though images for these markers were acquired from eight randomly selected non-overlapping fields under a 20× objective lens. For ACROSIN and DDX4 immunofluorescence, quantification differed: the integrated mean fluorescence intensity (MFI) of positive areas within each field was measured using ImageJ, background MFI was subtracted from the target region MFI to obtain corrected values, and the mean corrected MFI per field was calculated for each sample. Crucially, all images across all markers were captured using identical exposure times, laser intensities, and gain settings to ensure consistency.

### 2.7 Testis and brain proteomics analysis

Proteomic analysis of testicular and brain tissues from three control and three EMR-exposed mice was performed by Jingjie Biotechnology Company. Tissue proteins were extracted, and protein content was quantified using a BCA kit. Equal amounts of protein from each sample were subjected to trypsin digestion. Resulting tryptic peptides were dissolved in solvent A (0.1% formic acid in water) and loaded onto a homemade reversed-phase analytical column. Chromatographic separation was achieved on a Bruker Daltonics NanoElute UHPLC system at a flow rate of 500 nl/min using a binary mobile phase: solvent A (0.1% formic acid in water) and solvent B (0.1% formic acid in acetonitrile). Eluted peptides were analyzed online using a timsTOF Pro mass spectrometer in PASEF mode. Full MS scans (300–1500 m/z) were followed by MS/MS scans (400–850 m/z) with a 7 m/z isolation window, acquiring 20 PASEF MS/MS scans per cycle. DIA data were processed using DIA-NN (v.1.8). Tandem mass spectra were searched against the Mus musculus UniProt database concatenated with a reverse decoy database. Search parameters included: trypsin/P digestion with one missed cleavage allowed, fixed modifications, and variable modifications. Peptide spectral matches were filtered at a 1% false discovery rate (FDR).

### 2.8 Open field test (OFT)

The experiment is conducted in a quiet environment, where mice are randomly placed in the open field box (45cm×45cm×45cm) in sequence. Each time a mouse is placed, it is oriented towards a specific corner, with the central area serving as the middle zone. The mice are allowed to freely move for 5 mins, during which their movement trajectories are recorded using Anymaze software. Statistical indicators include: total distance, which measures the movement ability of mice; and central time ratio (time spent in center/total time), which measures the anxiety status of mice.

### 2.9 Tail Suspension Test (TST)

Gently remove the mice from their cages, apply adhesive tape 1 cm away from the tip of their tails, and attach the tails to the hook at the top of a tail suspension device measuring 55 cm in height, 60 cm in width, and 11.5 cm in depth, ensuring that the height between the tip of the animal’s tail and the ground is approximately 30 cm. Each mouse is videotaped for 6 mins, and analyzing the immobility time of the mice is an important indicator in the experiment, used to assess the degree of depression in the mice.

### 2.10 Y-maze

#### (1) Y-maze alternating behavior experiment

The Y-maze apparatus is made of three opaque plastic arms (labeled A, B, and C), with an angle of 120° between each arm. Mice are placed in the center of the maze and allowed to explore freely for 8 mins. The criterion for a mouse entering each arm is that all four limbs are fully inside. The mice enter three different arms in sequence (e.g., ABC, ACB, BCA, BAC, CAB, CBA), representing an alternation. The calculation method for the alternation score is: spontaneous alternation rate (%) = [(number of spontaneous alternations) / (total number of arm entries - 2)]×100. During the experimental interval between two mice, 75% alcohol is used to wipe the apparatus to avoid interference from odors.

#### (2) Measurement of spatial learning ability

Choose one arm as the “starting area” and another arm as the “novel area”. During the first test, place the animal with its back facing the center in the “starting area” while closing the “novel area”, allowing the animal to freely explore in both arms for 15 mins. After one hour, conduct the second test by placing the animal with its back facing the center in the “starting area” again while opening the “novel area”, allowing the animal to freely explore for 5 mins. The detection indicators include: the percentage of animals choosing the “novel area” as their preferred entry arm, the duration of stay in the “novel area”, and the number of entries and total arm entries for each arm.

### 2.11 Statistical analysis

GraphPad Prism 9 and R were used for all data analysis, graph plotting, and statistical analyses. Quantitative results were expressed as mean ± s.d. One-way analysis of variance (ANOVA), and two-tailed unpaired Student’s t-test were used to determine statistical significance. Exact P-values were provided accordingly in the figures or captions. P < 0.05 was used as the threshold for statistical significance; (*) indicates P < 0.05, (**) indicates P < 0.01, (***) indicates P < 0.001, and (****) indicates P < 0.0001. No statistical method was used to predetermine the sample size. The exact number of replicates and statistical tests are indicated in the figure legends. Unless otherwise indicated, n represents the number of independent experimental replicates.

## 3. Results

### 3.1 Sex-specific alterations in body weight and serum proteome following EMR exposure

Throughout the four-week exposure to 3.2 GHz electromagnetic pulse radiation, we measured food intake every five days and recorded body weight biweekly. No significant changes in food intake were observed in either male or female mice following EMR exposure (Figure 1A). However, body weight exhibited a sex-dependent response. While male mice showed no significant difference compared to controls, female mice displayed a marked decrease in body weight beginning in the third week of exposure (Figure 1B).

**Figure 1.**
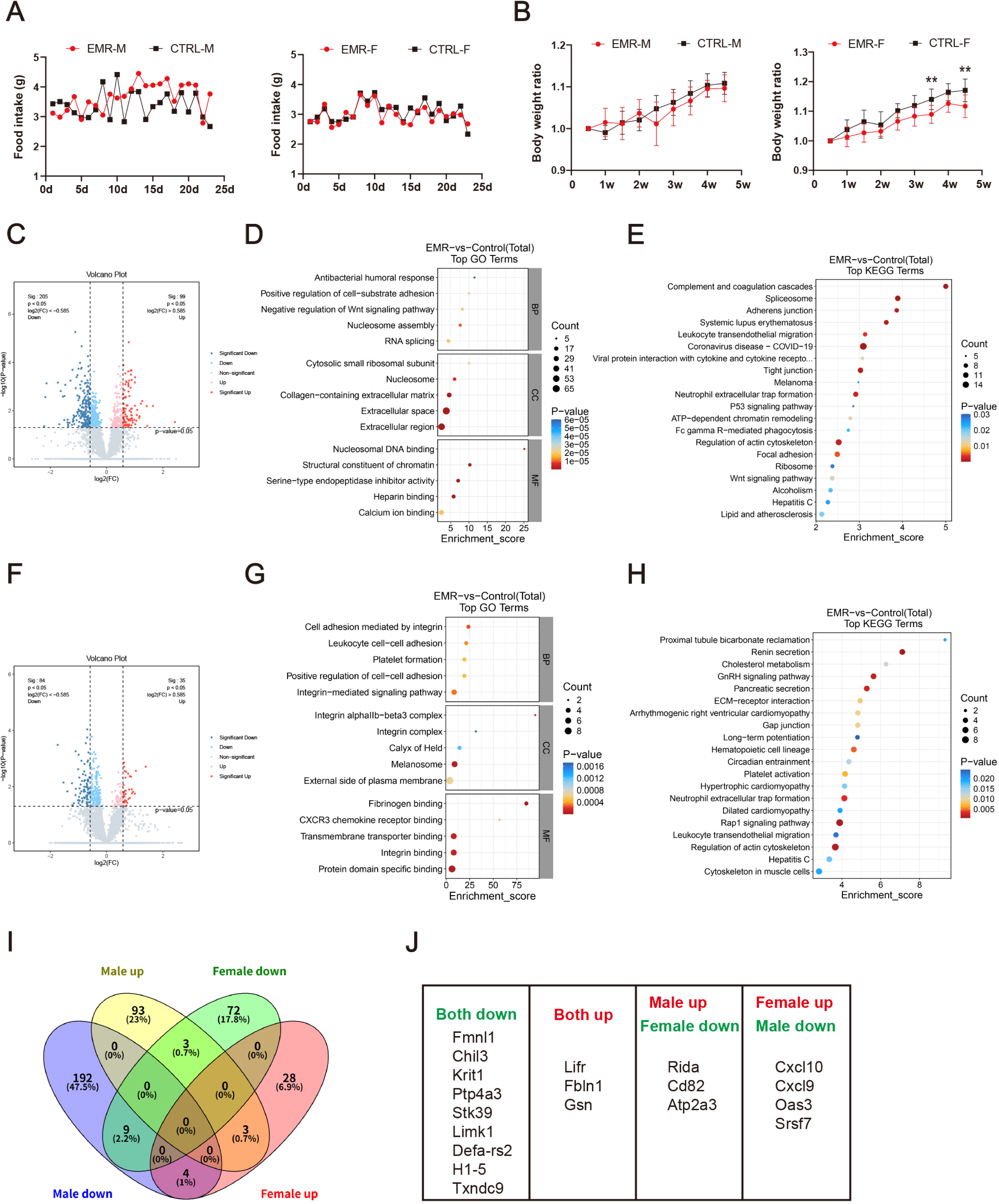
Sex-specific alterations in body weight and serum proteome following EMR exposure. (A) Dynamic changes in food intake of male and female mice; (B) Dynamic changes in body weight ratio of male and female mice, body weight ratio represents the ratio of the weight at a certain point in time to the initial weight; (C) Volcano plot of serum proteomic profiling of male mice. (D) GO functional enrichment analysis of DEPs in the serum of male mice. (E) KEGG pathway analysis of DEPs in the serum of male mice. (F) Volcano plot of serum proteomic profiling of female mice. (G) GO functional enrichment analysis of DEPs in the serum of female mice. (H) KEGG pathway analysis of DEPs in the serum of female mice. (I) Venn diagram of DEPs in serum of male and female mice. (J) List of commonly different proteins in serum of male and female mice.

To further investigate systemic responses, serum proteomic profiling was conducted. In male mice, 304 differentially expressed proteins (DEPs) were identified (205 down-regulated, 99 up-regulated; fold change ≥ 1.5 or ≤ 0.667, P < 0.05) (Figure 1C). Gene Ontology (GO) enrichment analysis revealed that these DEPs were primarily involved in antibacterial humoral response, positive regulation of cell-substrate adhesion, negative regulation of the Wnt signaling pathway, and nucleosome assembly (Figure 1D). KEGG pathway analysis showed significant enrichment in complement and coagulation cascades, spliceosome, and adherens junction pathways (Figure 1E).

In female mice, 119 DEPs were identified (84 down-regulated, 35 up-regulated) (Figure 1F). GO terms significantly associated with these DEPs included cell adhesion mediated by integrin, leukocyte cell-cell adhesion, platelet formation, positive regulation of cell-cell adhesion, and integrin-mediated signaling pathway (Figure 1G). KEGG analysis indicated enrichment in pathways such as proximal tubule bicarbonate reclamation, renin secretion, cholesterol metabolism, GnRH signaling, and pancreatic secretion (Figure 1H). Comparative analysis highlighted both shared and sex-divergent proteomic alterations (Figure 1I). Nine proteins (Fmnl1, Chil3, Krit1, Ptp4a3, Stk39, Limk1, Defa-rs2, H1-5, Txndc9) were down-regulated in both sexes, while three (Lifr, Fbln1, Gsn) were up-regulated in both. Notably, several proteins exhibited opposite expression trends: Rida, Cd82, and Atp2a3 were up-regulated in males but down-regulated in females, whereas Cxcl10, Cxcl9, Oas3, and Srsf7 were up-regulated in females and down-regulated in males (Figure 1J). These results demonstrate distinct sex-specific alterations in the serum proteome following EMR exposure.

### 3.2 EMR exposure induces male-specific testicular dysfunction and spermatogenic failure in mice

In female mice, four weeks of exposure to 3.2 GHz electromagnetic pulse radiation did not induce significant disruption of the oestrous cycle. Mice in the EMR group exhibited normal oestrus stages (Figure 2A), and ovarian histology revealed the presence of antral follicles and corpora lutea (Figure 2B), indicating preserved ovulatory function. Consistent with this, TUNEL staining and Ki67 immunohistochemical analysis showed no significant changes in ovarian apoptosis or cell proliferation levels following EMR exposure (Figure 2C, D).

**Figure 2.**
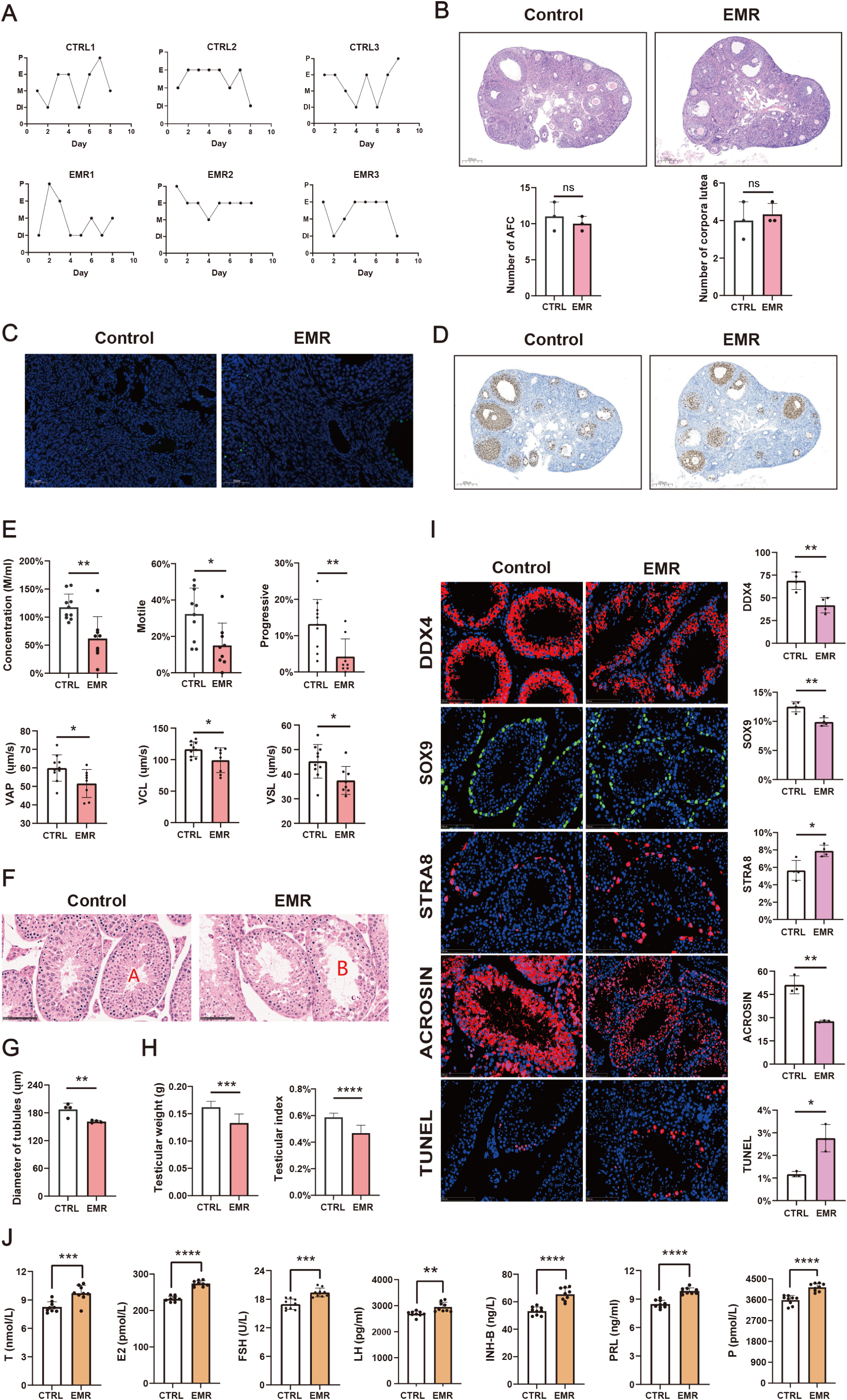
EMR exposure induces male-specific testicular dysfunction and spermatogenic failure in mice. (A) Changes in estrus cycle of female mice over 8 consecutive days. (B) Representative images of HE staining of female mouse ovaries after EMR exposure (Scale bar = 500 µm), and the count of antral follicles and corpora lutea in the ovaries (n = 3); (C) TUNEL staining of female mouse ovaries after EMR exposure (Scale bar = 200 µm); (D) Ki67 staining of female mouse ovaries after EMR exposure (Scale bar = 500 µm). (E) CASA experiment detects sperm parameters after EMR exposure; (F) Representative images of testicular HE staining magnified 40×; (G) Statistical results of the average diameter of 20 randomly selected seminiferous tubules from each section; (H) Testicular weight and testicular index (the ratio of testicular weight and body weight); (I) Representative images and fluorescence analysis of DDX4, SOX9, STRA8 and ACROSIN immunofluorescence staining, and TUNEL staining of testicular tissue after EMR exposure, Scale bars = 100 μm, Images from each section and each region of interest were captured at 40× magnification. (J) Reproductive hormone levels of male mice before and after EMR exposure; T:testosterone, E2: estradiol, INH-B: inhibin B, PRL: prolactin, P: progesterone.

In contrast, male mice displayed pronounced reproductive impairment after the same exposure regimen. Computer-assisted sperm analysis (CASA) revealed a significant reduction in sperm concentration, total motility, and the percentage of progressively motile sperm. Key kinematic parameters, including straight-line velocity (VSL), curvilinear velocity (VCL), and average path velocity (VAP), were also markedly decreased (Figure 2E). Histopathological examination showed notable alterations, featuring reduced seminiferous tubule diameter, widened intercellular spaces among spermatogenic cells, disorganized spermatogenic epithelium, and frequent vacuolization within the tubules (Figure 2F, G). Testicular weight and the testis-to-body weight ratio were significantly lower in the irradiated group (Figure 2H). Further immunofluorescence analysis of testes confirmed a significant decline in the number of spermatogonia (DDX4 +), Sertoli cells (SOX9 +), and mature sperm (ACROSIN +). In contrast, the population of unactivated spermatogonial stem cells (STRA8 +) increased significantly. TUNEL staining indicated elevated apoptosis within the testicular tissue (Figure 2I). Accompanying these morphological and cellular changes, the serum concentrations of reproductive hormones, including FSH, LH, estradiol, testosterone, inhibin B, progesterone, and prolactin, were significantly elevated in EMR-exposed males (Figure 2J).

Collectively, four weeks of 3.2 GHz electromagnetic pulse radiation exposure substantially impaired spermatogenesis, testicular structure, and endocrine balance in male mice, whereas female reproductive cyclicity, ovarian histology, and cellular homeostasis remained comparatively unaffected.

### 3.3 Identification of serum biomarkers for EMR-induced testicular dysfunction in male mice

To elucidate the molecular alterations underlying testicular damage induced by 3.2 GHz EMR, we performed proteomic analysis on testicular tissue. Principal component analysis (PCA) revealed substantial EMR-induced alterations in the testicular proteome (Figure 3A). Using a threshold of fold change ≥1.5 or ≤1/1.5 with P < 0.05, we identified 102 significantly downregulated and 99 upregulated proteins in irradiated testes (Figure 3B). GO enrichment analysis indicated that the downregulated proteins were predominantly associated with spermatogenesis and sperm function, including pathways such as “spermatogenesis,” “flagellated sperm motility,” and “sperm fibrillar sheath” (Figure 3C). Conversely, upregulated proteins were primarily enriched in pathways related to DNA damage repair (Figure 3D). Representative expression patterns of key spermatogenesis-related proteins are shown in Figures 3E and F.

**Figure 3.**
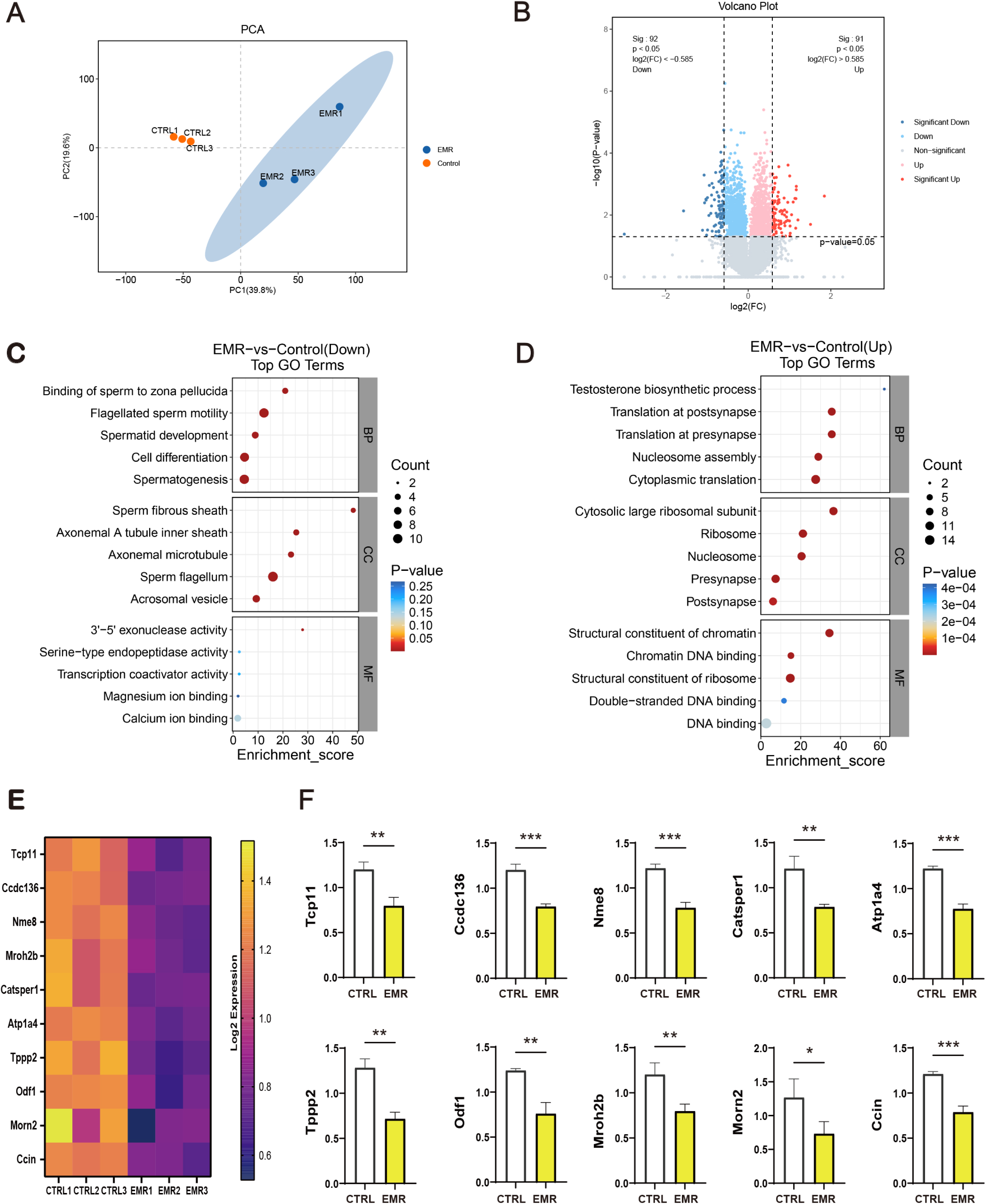
Proteomic analysis of testicular tissue after EMR exposure at 3.2 GHz for four weeks. (A) Principal component analysis (PCA) plots; (B) Volcano plot of expressed differential proteins after EMR-exposure; (C) GO functional enrichment analysis of downregulated proteins; (D) GO functional enrichment analysis of upregulated proteins; (E) Heat map of protein expression levels related to spermatogenesis; (F) Statistics of FPKM values for protein expression levels related to spermatogenesis.

Intersection analysis of differentially expressed proteins in testis and serum pinpointed a panel of seven proteins—Mdk, Rhoc, Ncf2, Hp1bp3, H1-1, H1-4, and H1-5—whose expression was consistently and significantly altered in both compartments (Figure 4A, B). Pearson correlation analysis demonstrated that the serum levels of these proteins were strongly and positively correlated with key sperm quality parameters, including sperm concentration, motility, and the percentage of progressive motile sperm (Figure 4C). Collectively, these proteins represent promising candidate serum biomarkers for predicting EMR-induced testicular impairment and diminished sperm quality.

**Figure 4.**
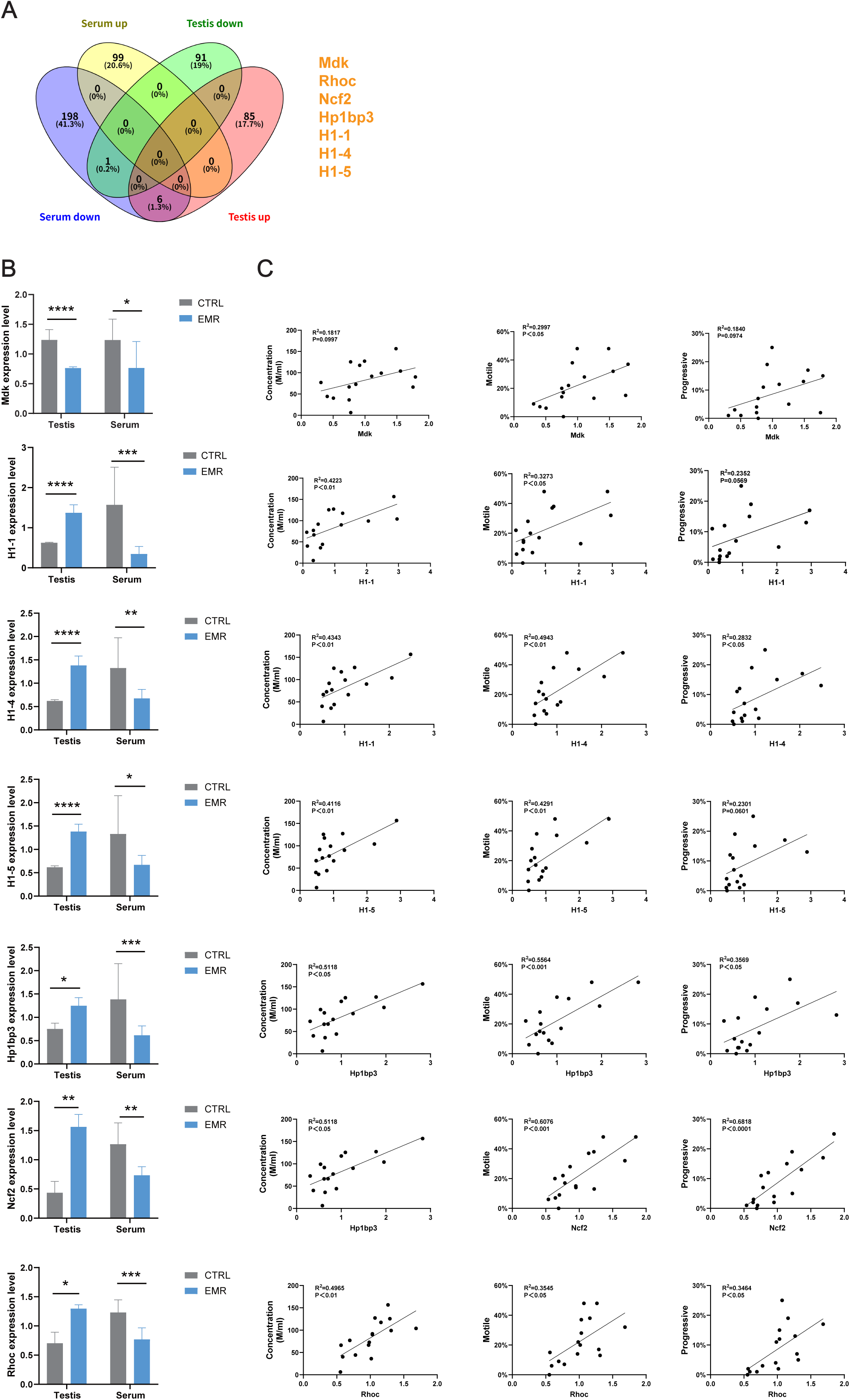
Serum indicators predicting reproductive impairment in male mice caused by EMR exposure. (A) Venn diagram of serum differential proteins and testicular differential proteins of male mice after EMR exposure; (B) Bar charts depicting the expression levels of commonly differentially expressed proteins in serum and testis; (C) Pearson correlation regression analysis of serum expression levels of common differentially expressed proteins and sperm parameters.

### 3.4 EMR exposure induces sex-specific neurobehavioral deficits in female mice

Behavioral assessments were conducted to evaluate the impact of four-week EMR exposure on mood and cognitive function. In the tail suspension test, while no significant changes were observed in male mice, EMR-exposed females exhibited a significant decrease in struggle time and a concomitant increase in immobility time compared to controls, indicating a depressive-like phenotype (Figure 5A). In the open field test, although statistical significance was not reached, both exposed sexes showed a trend toward increased total distance traveled and a decreased proportion of movement in the center zone, suggesting a potential anxiety-like response (Figure 5B). The Y-maze test, which assesses spatial working and reference memory, revealed that short-term spatial learning (spontaneous alternation) was unaffected in both sexes. However, in the long-term memory test, exposed female mice showed a significantly reduced novel arm alternation rate. Intriguingly, these females also spent more time and traveled a greater distance in the novel arm, indicating possible deficits in environmental recognition or habituation (Figure 5C). These results demonstrate that EMR exposure preferentially induces emotional and cognitive disturbances in female mice.

**Figure 5.**
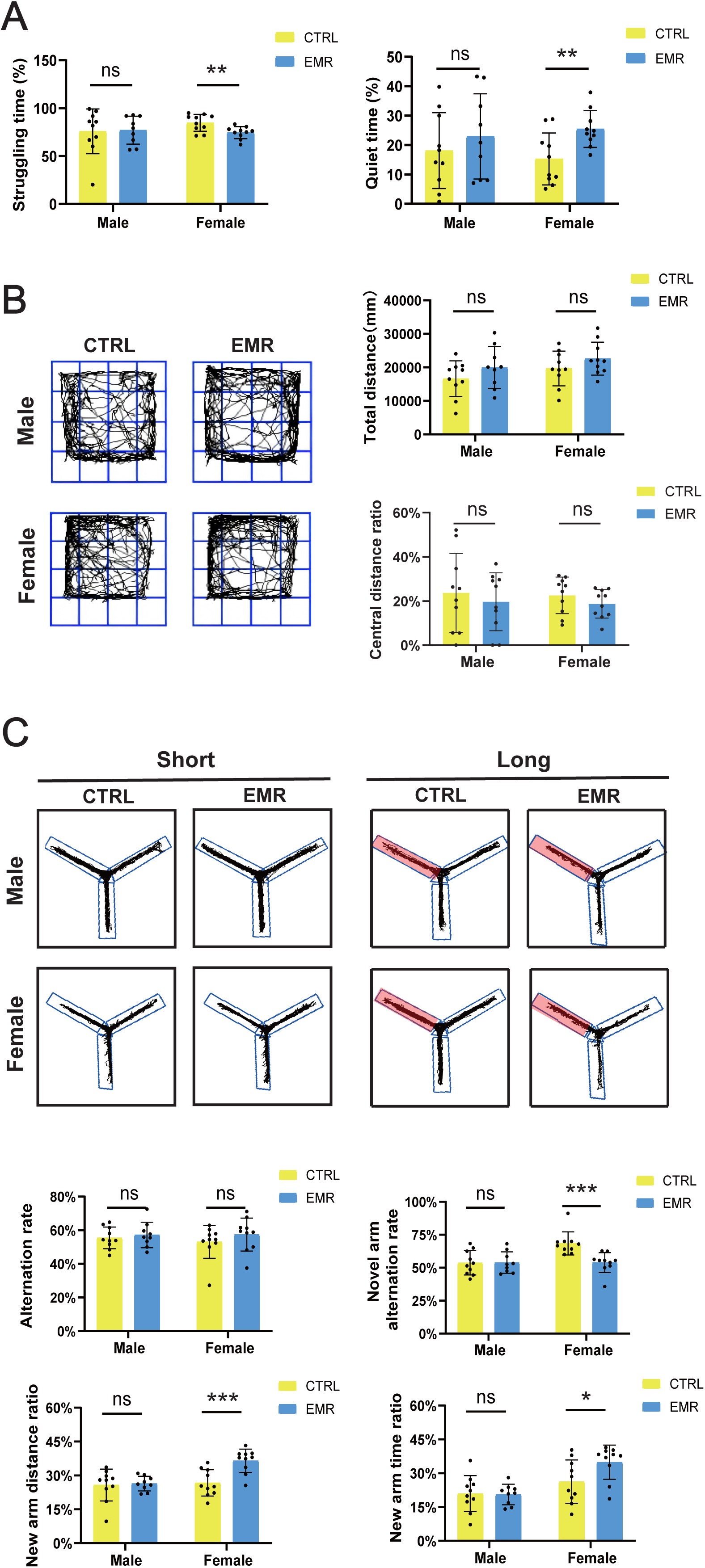
Behavioral analysis of male and female mice after EMR exposure at 3.2 GHz for four weeks. (A) The percentage of struggling and immobility time in the tail suspension experiment; (B) Representative trajectory diagram and index statistics of open field test; (C) Representative trajectory diagrams and indicator statistics of Y-maze alternation experiment and new arm experiment.

Given the central role of the hippocampus in emotion and cognition, we next examined its structural integrity. HE staining revealed a trend toward reduced thickness of the pyramidal/granule cell layers in the CA1, CA3, and DG subfields of both sexes post-EMR. This reduction reached statistical significance specifically in the CA1 region of female mice (Figure 6A, B). Furthermore, a marked increase in neurons with pyknotic nuclei was observed in the hippocampus of irradiated females but not males. Immunofluorescence staining for NeuN, a mature neuronal marker, confirmed a significant loss of NeuN-positive cells in the DG, CA1, and CA3 regions of EMR-exposed female mice (Figure 6C). This neuronal loss was accompanied by a significant increase in apoptotic cells, as detected by TUNEL assay, specifically in the female hippocampus (Figure 6D). These findings indicate that EMR inflicts selective hippocampal damage, characterized by neuronal loss and elevated apoptosis, with a pronounced female-specific vulnerability.

**Figure 6.**
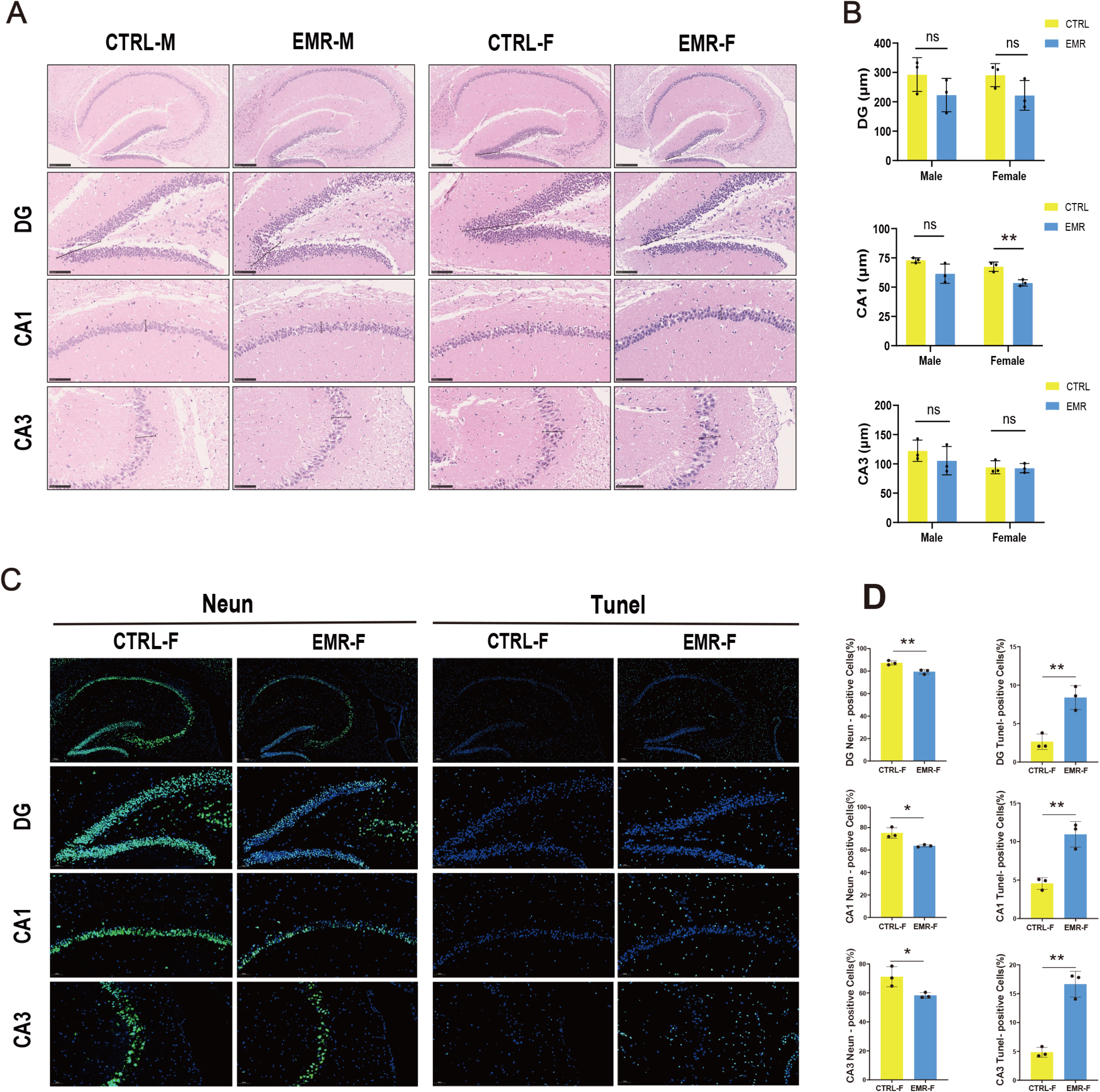
Hippocampal neuronal degeneration in mice after EMR exposure at 3.2 GHz for four weeks. (A) HE staining showing somatosensory cortex and different subregions of hippocampus (CA1, CA3, and DG); (B) Measurement of pyramidal nerve cell thickness in different areas of the corpus cavernosum after EMR-exposure; (C) Representative photomicrographs of NeuN-stained sections of female control and EMR-exposed mice (Scale bar = 50 µm); (D) Statistical comparison of the number of NeuN positive cells in the hippocampal subregions with respect to control.

### 3.5 KIF13A as a potential serum biomarker for EMR-induced neurological Impairment

To explore the molecular basis of EMR-induced neurotoxicity, we conducted proteomic sequencing on brain tissue from female mice. PCA confirmed distinct proteomic changes following exposure (Figure 7A). We identified 21 upregulated and 21 downregulated proteins (FC ≥1.5 or ≤1/1.5, P < 0.05) in the brain (Figure 7B). GO analysis showed upregulated proteins were involved in processes like protein tyrosine kinase activity regulation, while downregulated proteins were enriched in pathways critical for neurodevelopment and axon guidance, including “forebrain neuron differentiation” and “cerebral cortex neuron differentiation” (Figure 7C, D). Key neurogenesis-related genes (Gas6, Slit1, Rac3, Gdpd2) were notably downregulated (Figure 7E). Guided by the proteomic suggestion of prefrontal cortex (PFC) involvement, immunofluorescence confirmed significant neuronal loss (reduced NeuN+ cells) and increased apoptosis in the PFC of exposed mice (Figure 7F).

**Figure 7.**
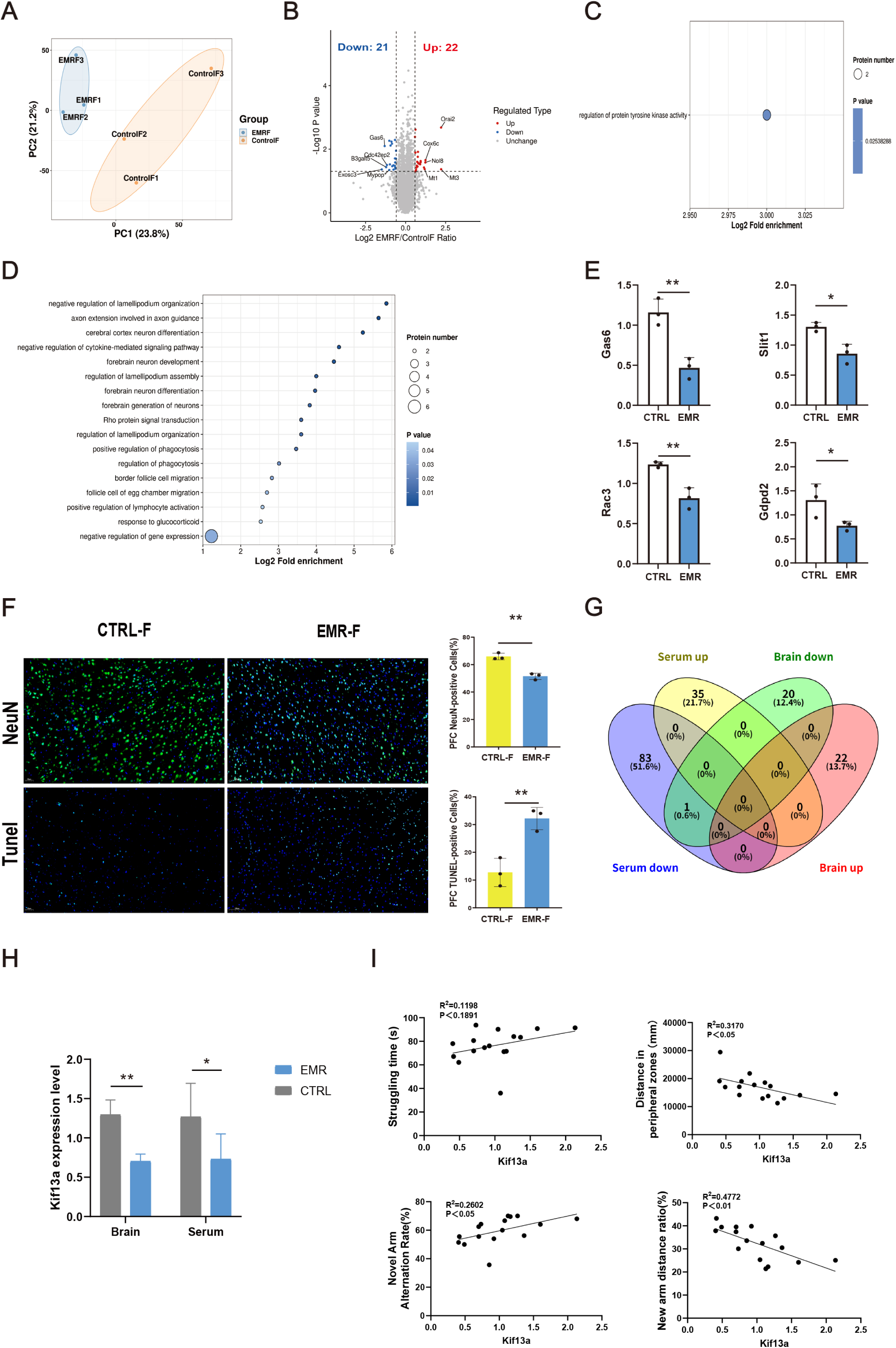
KIF13A may serve as a serum biomarker for predicting brain damage induced by EMR. (A) Volcano plot of differentially expressed proteins in brain proteomics sequencing after EMR exposure in female mice; (B) PCA plots of the brain proteomics sequencing; (C) GO functional enrichment analysis of upregulated proteins. (D) GO functional enrichment analysis of downregulated proteins; (E) Statistics of FPKM values for protein expression levels related to neuronal generation and development; (F) Representative images of NeuN and TUNEL staining of PFC, and statistical analysis of the number of positive cells (Scale bar = 100 µm); (G) Venn diagram of serum differential proteins and brain differential proteins of female mice after EMR exposure; (H) Bar charts depicting the expression levels of Kif13a in serum and brain; (I) Pearson correlation regression analysis of serum expression levels of Kif13a and behaviour test parameters.

Intersection analysis revealed a significant reduction in kinesin family member 13A (KIF13A) in both serum and brain tissue, as shown in Figures 7G and 7H. Correlation analysis demonstrated that serum KIF13A levels were positively associated with behavioral indicators of resilience, including struggle time in the tail suspension test, and with memory performance measured by alternation rate in the Y-maze. Conversely, KIF13A levels were negatively correlated with anxiety-like behavior, such as activity in the periphery of the open field (Figure 7I). These findings collectively identify serum KIF13A as a promising biomarker for emotional dysfunction and cognitive impairment associated with EMR exposure.

## 4. Discussion

The escalating ubiquity of radiofrequency electromagnetic radiation has positioned it as a significant environmental factor with considerable public health implications. Accumulating evidence suggests that the biological effects of such radiation may not be uniform across sexes, with emerging studies pointing to systematic differences in molecular pathways, target organ sensitivity, and physiological outcomes between males and females[20, 21]. In this study, we conducted a comprehensive, sex-stratified investigation into the biological consequences of exposure to 3.2 GHz pulsed RF-EMR in mice. The results reveal a complex landscape of biological susceptibility, characterized by fundamentally distinct systemic, reproductive, and neurological responses between males and females.

Our initial observations identified divergent systemic adaptations to EMR stress. A progressive decline in body weight among female mice from the third week of exposure, occurring without any alteration in food intake, points towards a sex-specific metabolic perturbation not seen in males. This physiological divergence was further reflected at the molecular level through profound differences in the serum proteome. While a limited set of proteins demonstrated common alterations across both sexes, the majority of differentially expressed proteins were uniquely modulated in one sex or exhibited directly opposing expression trends. Subsequent pathway analysis indicated that the male systemic response involved processes such as complement activation and cellular adhesion dynamics, consistent with previous reports of EMR-induced inflammatory and structural responses [22, 23]. In contrast, the female response was directed towards pathways regulating renal function, cholesterol homeostasis, and notably, the gonadotropin-releasing hormone signaling cascade, a key regulator of the reproductive axis that may underlie sex-specific endocrine disruption [13, 24]. These findings collectively establish that EMR exposure initiates two separate and distinct systemic physiological programs from the outset, thereby creating the foundational context for the organ-specific pathological vulnerabilities that were subsequently uncovered.

The investigation demonstrates a pronounced and specific male vulnerability localized to the reproductive axis. Exposure severely compromised the entire continuum of spermatogenesis, culminating in reduced testicular mass, a disorganized architecture of the seminiferous tubules, and a marked decline in all critical parameters of sperm quality, including concentration, motility, and kinematic properties. Converging insights from testicular and serum proteomics support a mechanistic model involving disruption at multiple stages of sperm production. Firstly, the observed significant reduction in midkine (Mdk) provides a mechanistic explanation for two key phenotypic changes. Mdk is a growth factor essential for maintaining spermatogonial stem cells in a proliferative, undifferentiated state[25]. Its downregulation likely undermines this maintenance, leading directly to the depletion of the germ cell reservoir, as evidenced by reduced DDX4+ cells, and, concurrently, to the aberrant activation of early differentiation signals, reflected in the increase of STRA8+ cells. Secondly, the persistent elevation of histone H1 variants alongside Hp1bp3 specifically within the testicular environment strongly suggests a direct impairment of the critical chromatin remodeling event where histones are replaced by protamines, a process indispensable for achieving the extreme DNA compaction required in mature sperm nuclei [26, 27]. Previous studies have demonstrated that disruptions in this histone-to-protamine transition can result in abnormal sperm morphology and compromised genetic integrity [28, 29]. The robust correlation between the serum levels of these candidate biomarkers, including Mdk and the histone H1 variants, and conventional sperm quality parameters further underscores their significant potential for the non-invasive assessment of RF-EMR-induced testicular injury.

In a stark and contrasting pattern, the primary site of female biological susceptibility was identified within the central nervous system. Exposed female mice exhibited a clear neurobehavioral phenotype encompassing depressive-like responses, anxiety-related behaviors, and specific deficits in long-term spatial memory. These functional impairments were consistently underpinned by tangible structural pathology, namely significant neurodegeneration and increased apoptotic activity within brain regions that are fundamental to affect and cognition, particularly the hippocampus and the prefrontal cortex. At the molecular level, our analysis identified a concurrent downregulation of the kinesin motor protein KIF13A in both cerebral tissue and systemic circulation. Given its well-established, essential role in the trafficking of synaptic receptors and its prior linkage to the regulation of anxiety [30–32] , the observed dysfunction of KIF13A represents a novel and plausible pathway mediating the neurological sequelae of EMR exposure specifically in females. Its measurable correlation with quantifiable behavioral metrics further nominates serum KIF13A as a promising and accessible biomarker for indicating RF-EMR-related neurobehavioral disruption.

## 5. Conclusion

This work delineates a clear and consequential sexual dimorphism in the organismal response to RF-EMR. Male susceptibility is predominantly concentrated within the reproductive axis, driven by a dual disruption targeting both the initial maintenance of the germline stem cell pool and the terminal maturation events of spermiogenesis. Conversely, female susceptibility is primarily neurological in nature, associated with synaptic transport dysfunction and selective vulnerability of limbic system structures. The identification of distinct, sex-specific circulating protein profiles offers a tangible and innovative foundation for the future development of precision biomarkers. We acknowledge certain limitations inherent to this study, including the application of a single carrier frequency and a continuous exposure paradigm, which may not fully encapsulate the complexity of real-world intermittent and multi-frequency exposure scenarios. Future investigations employing targeted genetic or pharmacological interventions will be essential to conclusively establish the causal relationships for the molecular mechanisms proposed here. Ultimately, these findings compellingly underscore the critical necessity of adopting a sex-aware framework in environmental health risk assessment and highlight specific molecular targets for the development of personalized preventive strategies against potential health risks associated with RF-EMR exposure.

## Acknowledgments

We are grateful to the Radiation Teaching and Research Office of the Naval Medical University for providing electromagnetic radiation equipment for this study.

## Author Statement

HY and KZ substantially contributed to the conception of the work. FL, ZL and CL contributed to the data analysis. KZ and FL wrote the manuscript. FL, JG, XY and JS helped to conduct animal experiments. HY and LW drafted and revised the manuscript. All authors read and approved the final manuscript.

## Data availability

Data will be made available on request.

---

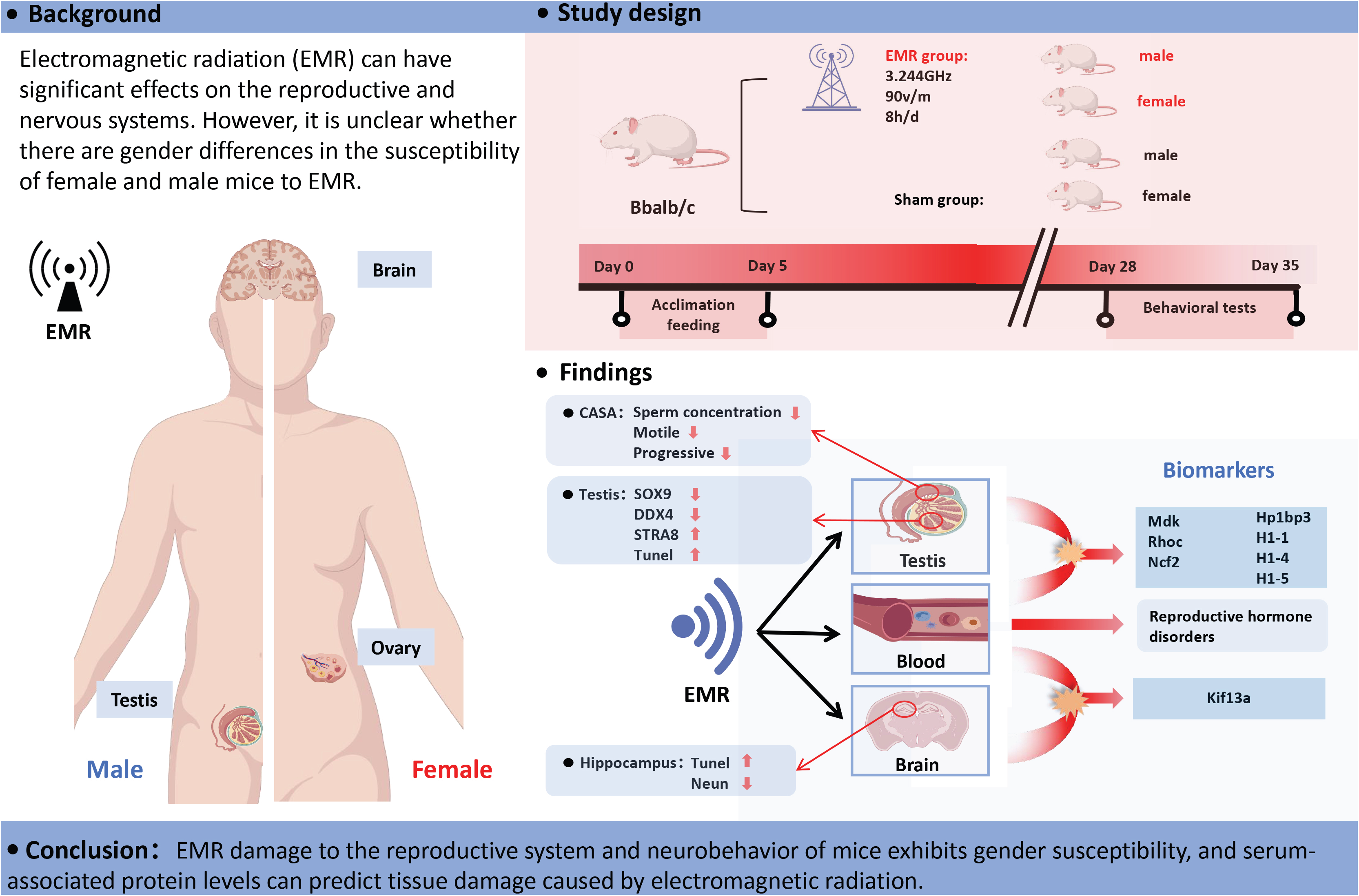

